# Primary Human Neutrophils and Monocytes/Macrophages Migrate along Endothelial Cell Boundaries to Optimize Search Efficiency

**DOI:** 10.1101/2024.06.24.600322

**Authors:** Nele Honig, Christina Teubert, Lucas Lamparter, Marius N. Keller, Judith Austermann, Philipp Berger, Anne Schmitz, Christiane Rasch, Harald Nüsse, Jürgen Klingauf, Luise Erpenbeck, Johannes Roth, Milos Galic

## Abstract

Neutrophils and monocytes/macrophages are sentinels of inflammatory signals. To reach the sites of action, both cell types attach to and then transmigrate the endothelial cell layer that lines the luminal side of blood vessels. While it has been reported that neutrophils and monocytes/macrophages actively migrate along the surface of the vasculature, it remains elusive if and how these motion pattern augment the efficiency of the immune system. Here, we conducted co-culture experiments of primary human monocytes and neutrophils, respectively, with human umbilical vein endothelial cells (HUVECs). Combining classical biomedical approaches with quantitative image analysis and numerical models, we find that immune cells simultaneously increase the number of sampled cells vs. traveled distance and sensitivity to chemokines by migrating along endothelial cell-cell boundaries. Collectively, these findings establish search optimization of neutrophils and monocytes/macrophages through limitation of motion pattern to cell-cell boundaries.

## Introduction

In response to an inflammatory signal, individual leukocytes display distinct responses. Monocytes participate in the clearance of pathogens, presentation of antigens, and aiding in tissue repair and regeneration. Complementary to monocytes, neutrophils migrate towards sites of infection, guided by chemical signals, where they neutralize pathogens. Collectively, their efforts contribute to pathogen protection and the maintenance of tissue health. While individual innate immune cells vary in their responses upon exposure to chemical cues, they all share the common challenge to detect the signal and depart from the blood stream. This process, called transmigration, relies on a finely orchestrated sequence of events and requires the availability of various surface receptors. Importantly, upon attachment monocytes and neutrophils both actively migrate towards future sites of transendothelial migration (TEM) ^1–4^. In addition, monocytes were shown to patrol the luminal site of the vasculature even in the absence of inflammation ^4–6^. However, whether any of these motion patterns along the surface of blood vessels is spatially confined, or if this may yield any benefits, is not known.

To address this aspect, we investigated the interactions of innate immune cells with endothelial cells that form the luminal surface of the vasculature. We find both immune cell types to migrate predominantly along cell-cell boundaries of the endothelial substrate. Numerical calculations suggest that the resulting motion patter present a hitherto unknown optimization of search efficiency.

## Results

### Primary human monocytes/macrophages migrate along cell-cell boundaries

In a first set of experiments, we aimed to probe the dynamics of primary human monocytes/macrophages on a confluent layer of primary human umbilical vein endothelial cells (HUVECs). To that end, primary monocytes/macrophages were added to a confluent HUVECs layer, and subsequently fixed. Consistent with previous reports ^4, 7^ only a subfraction of monocytes/macrophages did adhere to the endothelial sheet. Following, fixed samples were stained with the nuclear marker Hoechst and antibodies directed against *α*-Catenin, which links VE-Cadherin to the actin cytoskeleton ^8^. We find an enrichment of monocytes/macrophages at cell-cell boundaries of the confluent HUVEC layer (**Fig. 1a, top**).

**Figure 1.**
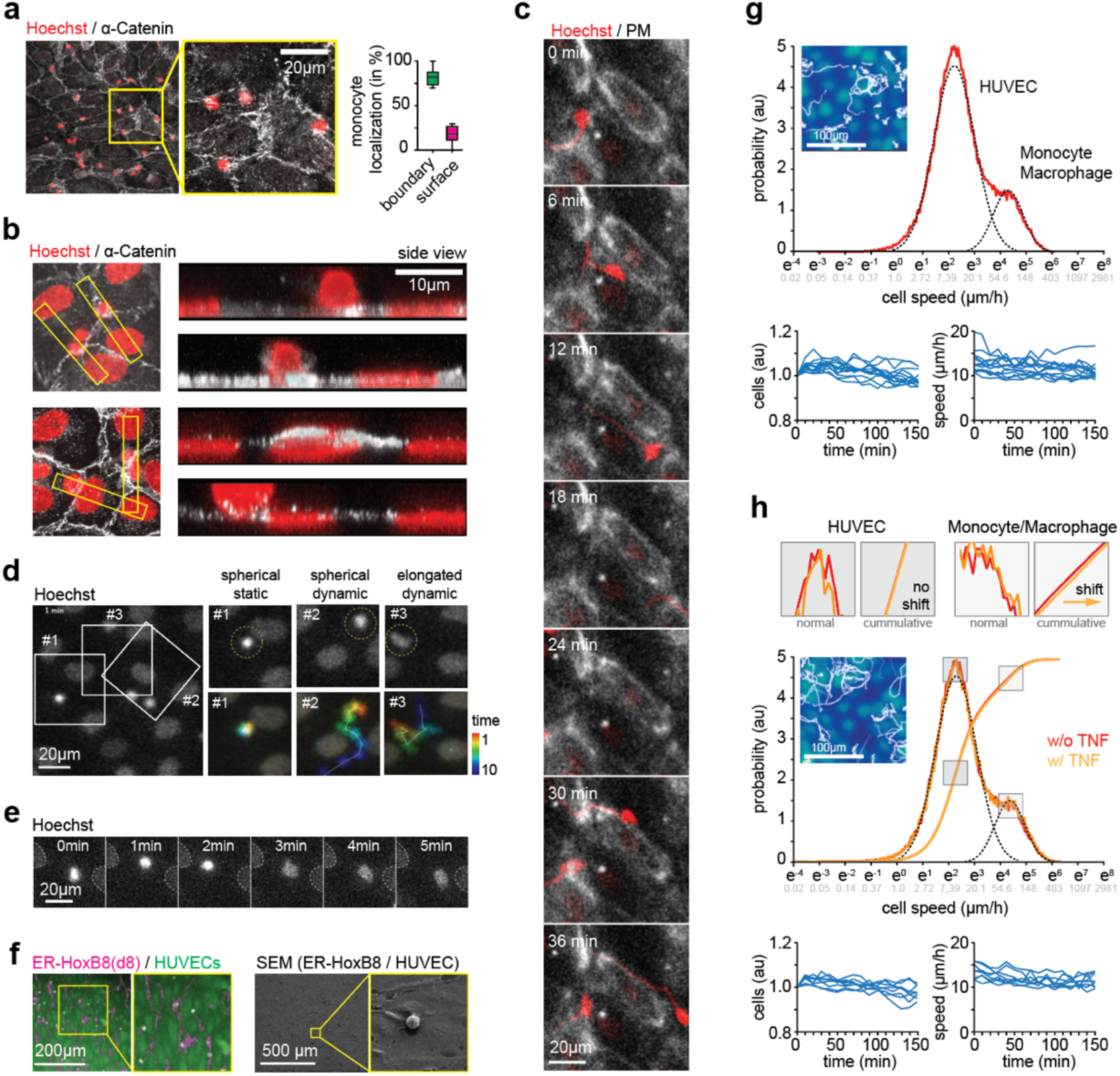
Primary human monocytes/macrophages cultured on top of monolayer of human umbilical vein endothelial cells migrate along cell-cell boundaries. **(a)** Primary human monocytes/macrophages (red) enrich at cell-cell boundaries of human umbilical vein endothelial cells (white). To the right, quantification of cells located at cell-cell boundaries (green) and on top of the cell (magenta) are shown (N = 3, n = 196 cells). **(b)** Immuno-cytochemistry indicates that primary human monocytes/macrophages transmigrate the HUVEC layer. **(c)** Primary human monocytes/macrophages (red) migrate along cell-cell boundaries (white) of human umbilical vein endothelial cells. **(d)** Time-lapse of primary human monocytes/macrophages (red) show different migration pattern. **(e)** Primary human monocytes/macrophages change fluorescence intensity of nucleus during transmigration. **(f)** ER-HoxB8 derived monocytes/macrophages migrate along HUVEC boundaries. To the left, HUVECs (green) and tracks (magenta) are shown. To the right, scanning electron microscope show that cells sit at cell-cell boundaries of a confluent HUVEC sheet. **(g)** Statistical analysis of motion pattern for primary human monocytes/macrophages and umbilical vein endothelial cells. At the top, representative tracks are shown. Below, quantification of cell dynamics is shown (N = 3, n = 12 technical repeats). **(g)** Migration of primary human monocytes/macrophages (magenta) and human umbilical vein endothelial cells in the presence of the pro-inflammatory signal TNFα. Again, motion pattern (top) and quantification (bottom) are shown (N = 3, n = 12 technical repeats). Scale bars, (a, c, d, e, g, h) 100 µm, (b) 10 µm, (g) 200 µm and 500 µm.

TEM is a multi-step progress, whereby cells tether, roll and adhere to endothelial cells, followed by transmigration from the luminal side of the blood vessel into the tissue ^9, 10^. To determine which of these steps were present in our system, we acquired 3D confocal sections of our co-culture system. We found monocytes/macrophages on top of a confluent HUVEC sheet, cells that were fixed while transmigrating through a junctional gap in the endothelial sheet, as well as cells located underneath the confluent HUVEC layer (**Fig. 1b**). Notably, the nucleus of monocytes/macrophages located on top of the layer appeared spherical, while the nucleus of cells underneath the endothelial sheet was flattened along the z-axis.

The experiments to this point show successful TEM of primary human monocytes/macrophages in our model system. The data, however, does not explain whether the high prevalence of monocytes/macrophages at cell-cell boundaries is caused by trapping or selective migration along these structures. To address this question, monocytes/macrophages and HUVECs were differentially labelled and simultaneously monitored using video-microscopy. Strikingly, we find monocytes/macrophages to travel selectively along cell-cell boundaries of HUVECs (**Fig. 1c, Fig. S1** and **Movie S1**). A detailed analysis of individual motion pattern identified three distinct motion patterns (**Fig. 1d** and **Movie S2**): Tracks that displayed no active motion, but moved at the same pace and in the same direction as the underlying HUVEC layer. Tracks that spent extended periods of time at one site and then actively moved to the next spot. And tracks that rapidly moved without extended dwell times at a particular site. While the former two groups were associated with round/bright nuclear staining, the latter one was associated with a dim/meandering nuclear staining (**Movie S3**). We find mutual spatial exclusion of the nucleus of monocytes/macrophages and HUVECs for all three motion types (**Fig. 1d**), indicative of an enrichment of monocytes/macrophages at cell-cell boundaries before, during and after TEM. Consistently, we find the nuclear staining to transition within less than one minute from bright/circular to dim/meandering shape (**Fig. 1e** and **Movie S4**). Unlike the transition from bright to dim, we did not observe any reverse transitions.

To further consolidate these observations, we took advantage of ER-HoxB8 cells that display a monocytes/macrophage-like phenotype upon induction ^11^. Much like the observations with primary human monocytes/macrophages, we find a striking aggregation of ER-HoxB8 derived monocytes/macrophages at cell-cell junctions of HUVECs, accompanied by migration primarily along these boundaries (**Fig. 1f**).

### Monocytes/macrophages and HUVEC dynamics in the healthy and inflamed state

The data to this point establishes dynamic migration of monocytes/macrophages along the surface of the vasculature in the absence of pro-inflammatory signals. While monocytes/macrophages extravasation at noninflamed sites is less common, these findings are consistent with previous reports ^2^. To systematically probe how a pro-inflammatory signal may affect the motion phenotype, we deployed three separate lines of investigation.

First, to set a base-line, we monitored cell dynamics of our co-culture system (**Movie S5**). To that end, cells were differentially labelled and monitored for 2.5 hours at an acquisition speed of 1 frame per minute. We find a bimodal distribution of cell speed, indicative of motion of HUVECs and monocytes/macrophages (**Fig. 1g**, top). Notably, we find no significant changes in cell number (**Fig. 1f**, bottom left) and cell speed (**Fig. 1g**, bottom right) across multiple technical and biological repeat, arguing for an overall healthy cell system throughout the whole imaging interval.

Second, we checked all time-lapse movies for spontaneous occurrences of cell death. For the ∼800 cells present in a field of view, we observed very few events over a period of 2.5 hours, further consolidating the over-all good health of the co-culture system. In the rare occasions where dissociation of the nucleus was observed (i.e. apoptosis), we monitored monocytes/macrophages dynamics prior to and during the event. Consistent with its described role, we find recruitment of monocytes/macrophages to dying cells, followed by local swarming and an uptake of cell debris (**Movie S6**). Strikingly, monocytes/macrophages did not migrate in a straight line, as a chemokine gradient would direct, but followed cell boundaries on their path to the source.

Finally, we exposed our co-culture system to TNFα, and probed for changes in cell dynamics (**Fig. 1h** and **Movie S7**). Again, cells were differentially labelled and monitored for 2.5 hours at an acquisition speed of 1 frame per minute. We find an increase in migration speed for monocytes/macrophages, but not for HUVECs. Again, motion pattern of immune cells remained restricted to cell-cell boundaries.

### Neutrophils also migrate along cell-cell boundaries

Having established that monocytes/macrophages preferentially migrate along cell-cell boundaries, we next aimed to probed the motion pattern of other innate immune cell type. Published work reported intraluminal crawling of neutrophils to distant emigration sites in wild-type mice ^1^. If this migration occurred in a particular form, however, remained elusive.

As above, we in a first set of experiments fixed primary human neutrophils cultured on HUVECs, and stained them with markers directed against the nucleus and cell-cell boundary. Similar to monocytes/macrophages, we find significant enrichment of primary human neutrophils at HUVEC cell-cell boundaries (**Figs. 2a,b**). Next, to quantify motion pattern, we monitored cell dynamics over 2.5 hours at a frame-rate of 1 image per minute. We find that neutrophils migrated as monocytes/macrophages along cell-cell boundaries (**Fig. 2c** and **Movie S8**).

**Figure 2.**
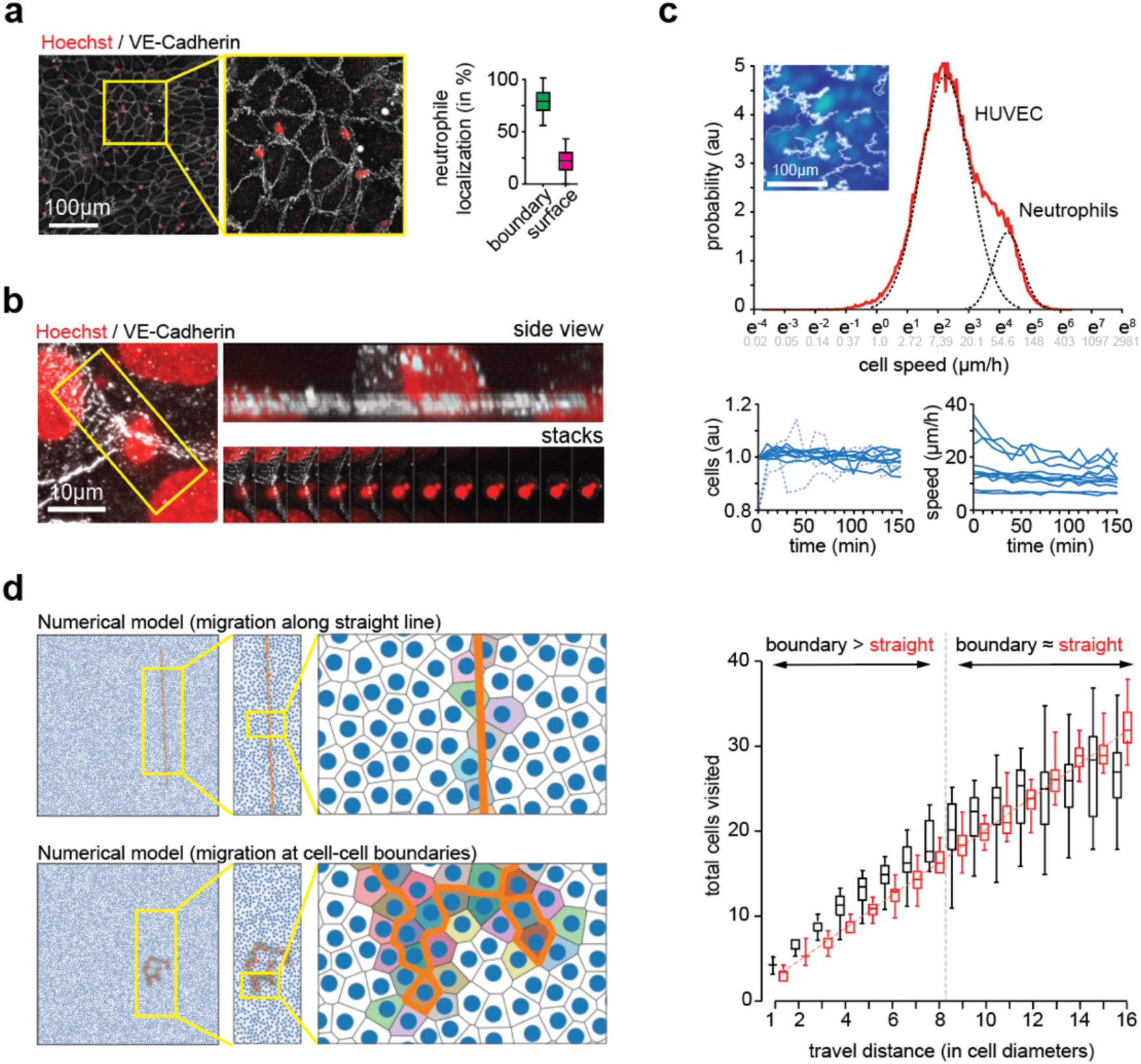
Primary human neutrophils migrate along cell-cell boundaries of human umbilical vein endothelial cells. **(a)** Primary human neutrophils cells (red) enrich at cell-cell boundaries of human umbilical vein endothelial cells (white). To the right, quantification of cells located at cell-cell boundaries (green) and on top of the cell (magenta) are shown (N = 3, n = 185 cells). **(b)** Primary human neutrophils cells migrate on top of human umbilical vein endothelial cells. **(c)** Statistical analysis of motion pattern of primary human neutrophils and umbilical vein endothelial cells. At the top, representative tracks are shown. Below, quantification of cell dynamics is shown (N = 3, n = 12 technical repeats). **(d)** Numerical model indicates that migration along cell-cell boundaries posits a better search strategy than a straight line for short distances. To the left, representative runs along straight line (top) and along cell-cell boundaries (bottom) are shown. To the right, a quantification of sampled endothelial cells vs. walking distances is shown for both motion types. Scale bars, (a, c) 100 µm, (b) 10 µm.

### Migration along cell-cell boundaries improves the search efficiency

Migration pattern were reported to influence the search efficiency of single cells ^12^. To address the question how migration along cell-cell boundaries may affect the search ability of immune cells, we derived a reductionist model that reconstitutes the basic motion pattern observed for neutrophils and monocytes/macrophages, respectively, on top of the vasculature. We then let individual cells migrated for a defined number of cell diameters, and scored the number of neighboring cells that were touched. For track lengths of less than 8 cell diameters, migration along the cell-cell boundaries allowed sampling more cells than a straight line (**Fig. 2d, Fig. S2** and **Movie S9**). For longer tracks, a straight line becomes more efficient in our numerical model. These findings are relevant, as it argues that at physiologically relevant length-scales restricting the motion of monocytes/macrophages to the cell-cell boundaries outperforms motion along a straight line, which is generally considered the optimal search strategy for planar system.

## Discussion

Monocytes were described to actively move from sites of adhesion to the nearest junction for diapedesis ^2^. However, a large fraction of monocytes will migrate multiple cell diameters along the luminal site of endothelial cells in vitro ^2^ and in vivo ^3^. Similarly, neutrophils will not pick the nearest available junction, but migrate along the vasculature to hotspots from which they depart into the tissue ^13^. Whether these ‘long walks’ along the endothelial layer are spatially restricted in a particular manner, or if such a bias may yield any benefits, has to date not been assessed. Here, we probed the motion pattern of primary human neutrophils and monocytes/macrophages on an endothelial layer. We find that monocytes and neutrophils, respectively, enrich at cell-cell boundaries of human umbilical vein endothelial cells, and dynamically move along these structures.

Conceptually, these findings yield two major conclusions. First, motion along cell-cell boundaries increases search efficiency. Neutrophils and monocytes/macrophages not only respond to diffusive, but also to membrane-bound signals. For instance, non-classical monocytes were described to sample the luminal site of the vasculature for unhealthy particles ^5, 6^. It is thus plausible to envision that some of the motion pattern observed at the luminal side of the vasculature may reflect patrolling of non-classical monocytes (**Fig. S3**). Similarly, neutrophils transmigration has been described to occur at hotspots ^13^. While fundamentally different objectives, both tasks will benefit by increasing the number of sampled endothelial cells per traveled distance. Noteworthy, the intercellular adhesion protein I-CAM appears to play a central role in the migration of monocytes ^2^ and neutrophils ^13^. Second, motion along cell-cell boundaries augments fidelity for diffusive chemokines. For an immune cell in the blood stream, the highest concentrations of soluble signaling cues that originates from a nearby inflamed tissue occurs at cell-cell junctions. From there, the signal intensity decreases quadratically with distance. Hence, to reliably detect a potential inflammation, it is beneficial to migrate along cell-cell boundaries.

Collectively, these findings establish how neutrophils and monocytes/macrophages increase in search efficiency without the need to improve migration speed or sensitivity to chemokines. The observation that all cells displayed migration along the cell-cell border strongly suggest that the observed phenotype is equally relevant to immune cell function in the inflamed and normal tissue. Future work will determine the relevance of these findings for other cell types.

## Materials and Methods

### Handling of human umbilical vein endothelial cells (HUVECs)

Commercially purchased HUVECs (Promocell, C-12208) were thawed, centrifuged at 400 x g and suspended in HUVEC Medium. HUVEC Medium consists of endothelial cell growth medium 2 (EGM2) (Promocell, C22011) and Medium 199 (Earle’s Salts, gibco, 31120022) with a ratio 1:1 and supplemented with 5 % fetal bovine serum (FBS) (Sigma Aldrich, S0615), 50 ng/µl amphotericin B (Gibco, 15290-018), 50 µg/ml gentamicin (Gibco, 15710-049), and 0.5 U/ml heparin (Sigma Aldrich, H3149). HUVECs were cultured in 6 cm and 10 cm CellBIND surface dishes (Corning, 3295 and 3296). HUVECs were no longer used after 2-3 passages. For passaging, cells were washed with 2 ml phosphate-buffered saline (PBS) (Gibco, 10010015) and detached by incubating with 1 ml trypsin/EDTA (0.04 % (w/v)/ 0.03 % (w/v), Promocell, C-41010) for three minutes at 37 °C. Cells were resuspended with 9 ml of culture medium and centrifuged at 400 x g for 4 minutes. The supernatant was discarded and cells were resuspended in HUVEC medium. The cells were then seeded into a new dish. Cells were passaged 1:3, each passage after 3 days. After two passages, once the cells had reached 90-100 % confluency, cells were frozen at -140°C in cryovials (Thermo Scientific, 144661), containing 90 % FBS and 10 % dimethyl sulfoxide (DMSO). 3 days after thawing the cells, they are seeded into ibidi µ-Slide VI that are coated with collagen. About 50,000 cells were seeded into each channel.

### Isolation and culturing of primary human monocytes/macrophages

Human monocytes/macrophages were purchased from the blood bank of the University Hospital Münster. In brief, monocytes/macrophages were isolated form thrombocones that were cut open at the bottom end and put in the opening of a T75-bottle. The top end of the thrombocones was also cut open and the blood was added to the T75 bottle, which was then filled up to 120 ml with HBSS. The mixture of blood and HBSS was then mixed by swaying the bottle. 30 ml of the mixture was then layered on 20 ml of pancoll. Following, cells were centrifugated at 540 x g for 35 minutes at room temperature. In the meantime, a second gradient was prepared by adding 15 ml of HBSS, 13.5 ml of Percoll and 1.5 ml of MEM 10* to centrifuge tubes. This mixture was centrifuged at 15,455 x g for 12 minutes, and then left in the fridge. After centrifugation of the HBSS-blood-Pancoll solution, a layer of monocytes formed in the center, which was carefully removed and added to a 50 ml falcon. The falcon was then filled with HBSS and centrifuged for 10 minutes at 300 x g. The supernatant was removed and the pellet was resuspended in 1 ml of HBSS and then filled up to 50 ml with HBSS. This solution was then subjected to two more rounds of centrifugation at 170 x g and 300 x g for 10 minutes each. After removing the supernatant and resuspending the pellet with 1 ml of HBSS, 1 ml was added to the tube containing the prepared second gradient. This tube was then centrifuged at 540 x g for 60 minutes. Afterwards, two layers should have formed. The top layer was carefully added to a new 50 ml falcon, which was then filled up to 50 ml with HBSS. This falcon was then centrifuged and filled three more times at 300 x g, 170 x g and 300 x g for 10 minutes each. The last pellet was resuspended with RPMI 1640 (Pan Biotech P04-17525), containing 15 % FCS (Biowest, S00GF1000J), 1 % Penicillin-Streptomycin (10,000 U/ml Pen,10mg/ml Strep, Pan Biotech, P06-07100), 1 % L-Glutamine (200mM, Pan Biotech, P04-80100), and 1 % non-essential amino acids. The purity of the monocytes was checked by FACS. Samples were used when purity > 80 %.

### Culturing and differentiation of ER-HoxB8 derived monocytes/macrophages

As previously published ^11^, ER-HoxB8 cells were cultured in RPMI (Promocell) containing 10 % FCS and 1 % Pen-Strep. The base medium was supplemented with 2 µg recombinant granulocyte-macrophage colony-stimulating factor (GM-CSF) (Immunotools), and 1 µm *β*-Estradiol (Sigma–Aldrich). Cells were passaged every other day in 6-well chambers. For differentiation, ER-HoxB8 cells were transferred into a 15 ml centrifuge tube and washed by centrifuging at 300 x g for seven minutes and resuspending in PBS. The falcon was then filled up to 12 ml with PBS and the process repeated twice. Finally, the cells were resuspended in 1 ml of the base medium, supplemented with GM-SCF but no estradiol. This counts as day 0 of differentiation.

### Isolation and culturing of primary human neutrophils

All experiments with human neutrophils were approved by the Ethics Committee of the University of Münster (Application #2021-657-f-S). Donors were fully informed about any possible risks and their right to withdrawal from the study at any time with informed consent confirmed in writing. Blood was drawn into S-Monovettes EDTA (7.5 mL, Sarstedt) and neutrophils were isolated from blood using the EasySep^TM^ direct human neutrophils isolation kit (Stemcell Technologies, 19666). They were then centrifuged at 400 x g for 10 minutes at room temperature, resuspended in RPMI medium and counted. Purity was determined to be over 98%.

### Live cell imaging

Monocytes/macrophages and neutrophils, respectively, were stained with Hoechst (1:200) and Cell Tracker Red (1:1000) and incubated in the dark at room temperature for 40 minutes. Neutrophils were centrifuged at 400 x g for 10 minutes, while monocytes/macrophages were centrifuged at 300 x g for 8 minutes and resuspended in HUVEC medium and centrifuged again to wash out the staining. The pellet was then resuspended with 200 µl of HUVEC medium per slide. The cells in the suspension were then counted and the suspension diluted to 1.5 x 10^6^ cells/ml. To seed neutrophils/monocytes, 30 µl of the cell suspension was added to the channel on one side and the HUVEC medium was removed by suction. This results in 45,000 cells in each channel of the ibidi µ-slide. The slide was left to rest for 1 h at 37 °C before channels were filled up with HUVEC medium and imaged. TNF*α* was added to cell medium at a concentration of 10 ng/ml, left to incubate overnight, and washed before seeding

### Immunocytochemistry

For antibody staining, neutrophils and monocytes, respectively, were seeded on HUVECs as described previously, skipping the staining with Hoechst and cell tracker. Instead, slides were washed with PBS++ and fixed with PFA 4 % dissolved in PBS++. Specifically, 30 µl of PFA was first added to one side of the channel, and removed from the other side. Then, 100 µl were added to each channel. PFA was left in the channels to incubate for 10 minutes at room temperature. The channels were then washed three times with PBS++. Following, fixed cells were washed with PBS++, and then permeabilized. The permeabilization solution was left in the channels to incubate for ten minutes at room temperature. The slides were then washed with PBS++. For blocking, 100 µl blocking solution was added to each channel and left to incubate on a shaker for 60 minutes at room temperature. Primary antibodies were diluted in PBST in a concentration of 1:200 and left to incubate overnight at 4°C on a shaker. Following primary antibodies were used: CD14 Monoclonal Antibody (Thermo Fisher, 60253-1-IG), VE-Cadherin Monoclonal Antibody (R&D Systems, MAB9381), *α*-Catenin (Atlas Antibodies, HPA063535). Secondary antibodies were diluted at a concentration of 1:1000 in PBST, and incubated in the dark at room temperature on a shaker for 60 minutes. Following secondary antibodies were used: Alexa fluor 488 goat-anti-mouse (Life Technologies, A11001), Alexa fluor 568 goat-anti-rabbit (Life Technologies, A11011). In some experiments, cells were alternatively stained with Hoechst (1:500) or Phalloidin (1:1000). Finally, channels were washed with PBS++. About three drops of ibidi mounting medium (ibidi, 50001) was added to the channels.

### Fluorescence microscopy

Cell migration was imaged using an inverted confocal microscope (Nikon, Eclipse Ts2) with a digital camera (Nikon DS-Fi2), using a 20x objective at 37°C and 5 % CO_2_ with a frame interval of 60 seconds. Using multi-frame imaging allowed the recording of multiple positions per channel (i.e. technical repeats). Fixed samples, which were stained using antibodies, were imaged using Leica fluorescence microscope. Z-Stacks were taken using a 63x objective. The distance between the single images in the z-stack was set to 0.5 µm.

### Scanning electron microscopy

Samples were washed three times in the chamber with PBS++, then fixed overnight using PFA (4%) and GA (2.5%) in PBS++. Next, the samples were washed three times for 5-10 minutes each in PBS++. The chamber bottom was then cut out to dimensions of approximately 3.8 x 10 mm. Subsequently, the samples were fixed for 1 hour with 1% OsO4 in PBS (without Ca and Mg). Following, an ascending alcohol series was applied (30%, 50%, 70%, 90%, followed by two rounds of 100% ethanol for 10-15 minutes each), followed by critical point drying using a Leica CPD300 automatic system. Finally, samples were mounted using Carbon Adhesive Tabs (Plano G3347) and sputter-coated with 15 nm of Au. Imaging was conducted with a scanning electron microscope FEI Quanta 600F at 15 kV.

### Image analysis

For tracking, movies were first pre-processed in Fiji/ImageJ ^14^ as follows: first contrast enhanced, using 0.35% saturated pixel (equalize histogram, process all), followed by image alignment via SIFT ^15^, using following parameter conditions: initial gaussian blur = 1.6 pix, 3 steps per octave, minimum image size: 64 pix, maximum image size: 1024 pix, feature description size: 4, feature descriptor orientation bins 8, closest/next ratio: 0.92, max alignment error: 25 pix, inlier ratio: 0.05, expected transformation: translation, output: interpolate. Next, individual nuclei were identified using Stardist ^16^ with following parameters: Model: versatile (fluorescent nuclei); normalized image; percentile low: 1%, percentile high: 100.0%, probability/score threshold: 0.40, overlap threshold: 0.40, output type: both, ROI position: Automatic. The output was analyzed with Trackmate ^17^, using the following parameters: LoG detector, object diameter: 25, quality threshold: 0.03, simple LAP tracker, linking distance: 3, gap-closing distance: 3, gap-closing max frame gap: 30. Tracks were displayed tracks backwards in time for 30 frames.

### Numerical model

To investigate the influence of migration patterns on the search efficiency of immune cells, a computational model implemented in Python was developed (source code will be deposited on GitHub upon acceptance). The code models immune cell migration along cell-cell boundaries and as a straight line, thus allowing the analysis of migration strategies. The user can customize parameters such as cell radius, number of simulated runs, length of the migration path, and visualization options. Upon execution, the code generates images as JPEG files of the migration paths and provides a statistical analysis using the Mann-Whitney U test to compare the distribution of visited regions between simulated migration paths at cell-cell boundaries and as straight lines. The results were visualized using box plots and saved with the generated JPEGs of the migration pathways in a separate folder.

### Statistics

Unless stated otherwise, a non-parametric rank-sum test was used to determine significance (ns for p> 0.05; * for p<0.05; ** for p<0.01; *** for p<0.001). Box plots present the median, extending from the 25^th^ to the 75^th^ percentile. Whiskers depict to the smallest and largest values.

## Acknowledgments

We would like to thank Jenny Lücking for excellent technical assistance, and the members of the Galic lab for critical reading of the manuscript. MG was supported by funding from the DFG (GA2268/3-1, GA2268/4-1, CRC1450 N07). JR was supported by grants of the Interdisciplinary Center of Clinical Research at the University of Munster (Ro2/007/22), and the DFG (CRU 342 P3, TRR332 B5, CRC1450 C1). LE received support from the DFG (TRR332) and the Interdisciplinary Center of Clinical Research at the University of Munster (Erp2/016/23). JK acknowledges funding from the DFG (CRC 1348 A02). NH and MNK received funding from the MedK program of the Medical Faculty of the University of Münster.

## Author contributions

NH and CT prepared the co-culture system and acquired all data. LL, MNK and MG prepared the numerical model. JA, PB and JR assisted with monocytes/macrophages. AS and LE assisted with neutrophils. CR, HN and JK helped with SEM imaging. MG analyzed data and wrote the manuscript with input from all authors.

## Data availability

Code will be deposited online on our homepage and GitHub upon acceptance. All raw data will be made available upon request.

## Supporting Information for

This PDF file includes:

Figures S1 to S3

Legends for Movies S1 to S9

### SUPPLEMENTARY FIGURES

**Figure S1.**
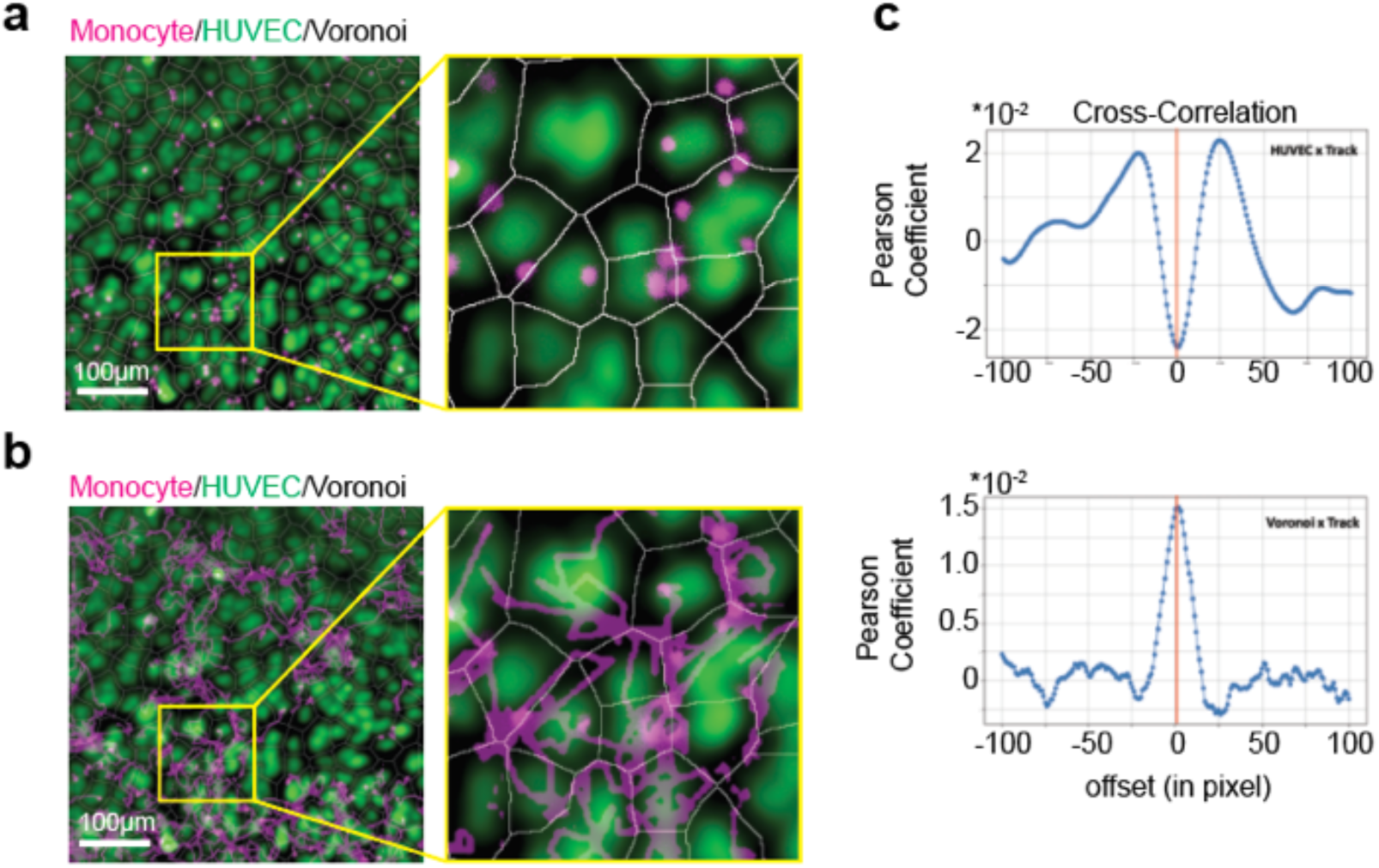
Validation of primary human monocytes co-cultured with primary human endothelial cells. **(a)** Monocytes (magenta) locate to Voronoi (white) derived from HUVEC signal (green). **(b)** Representative image of cells migrating along cell-cell boundaries. **(c)** Cross-cor relation analysis showing mutual exclusion of monocyte vs. HUVEC (top) as well as co-enrichment of monocytes vs. voronoi (bottom). (N = 8, n_HUVEC/voronoi_ = 2276 cells, n_monocytes_ = 1667 cells). Scale bars, (a, b) 100 µm.

**Figure S2:**
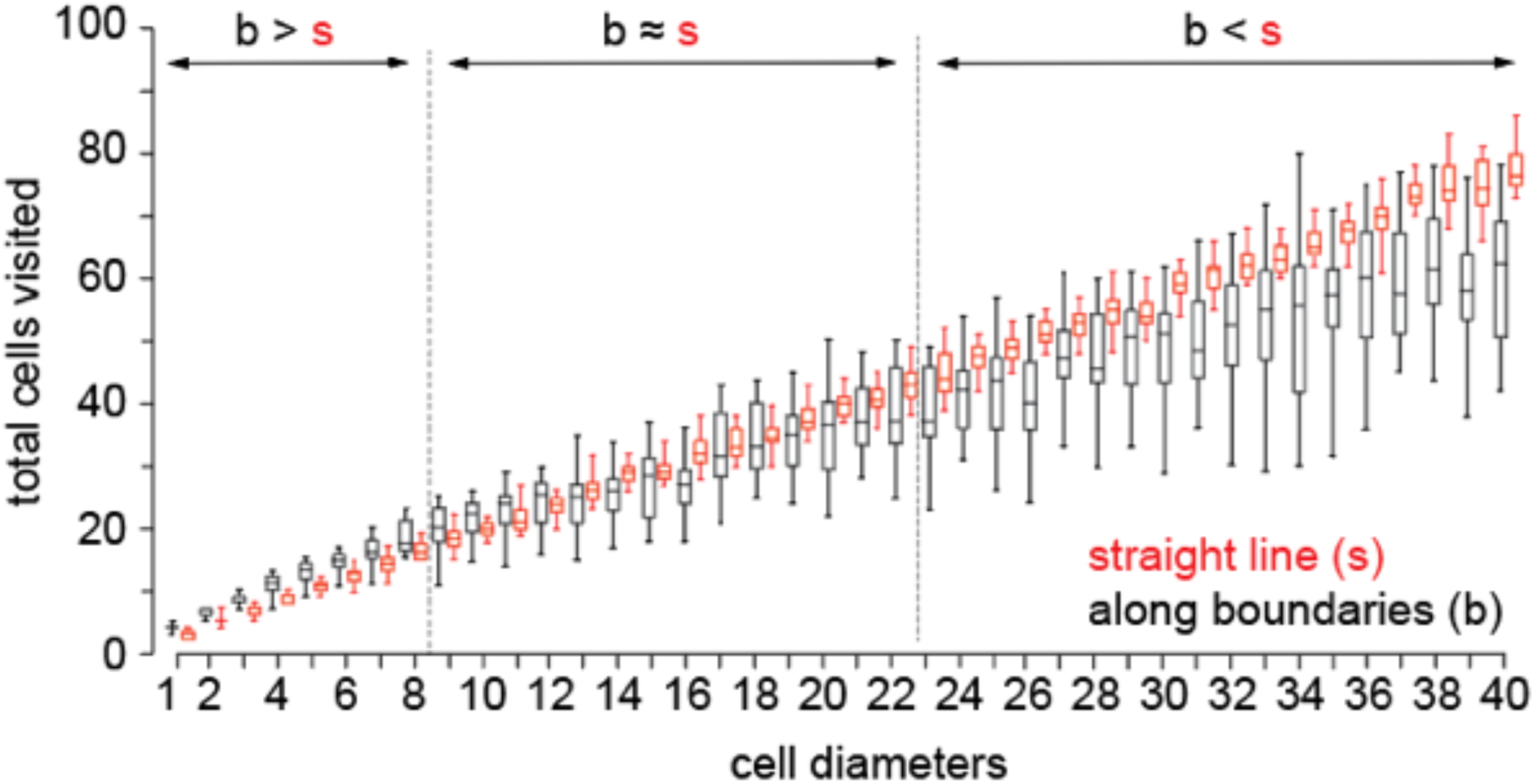
Numerical model indicates that migration along cell-cell boundaries is a better search strategy for short distances, while a straight line is better for distances. Quantification of cells encountered for different walking distances is shown.

**Figure S3.**
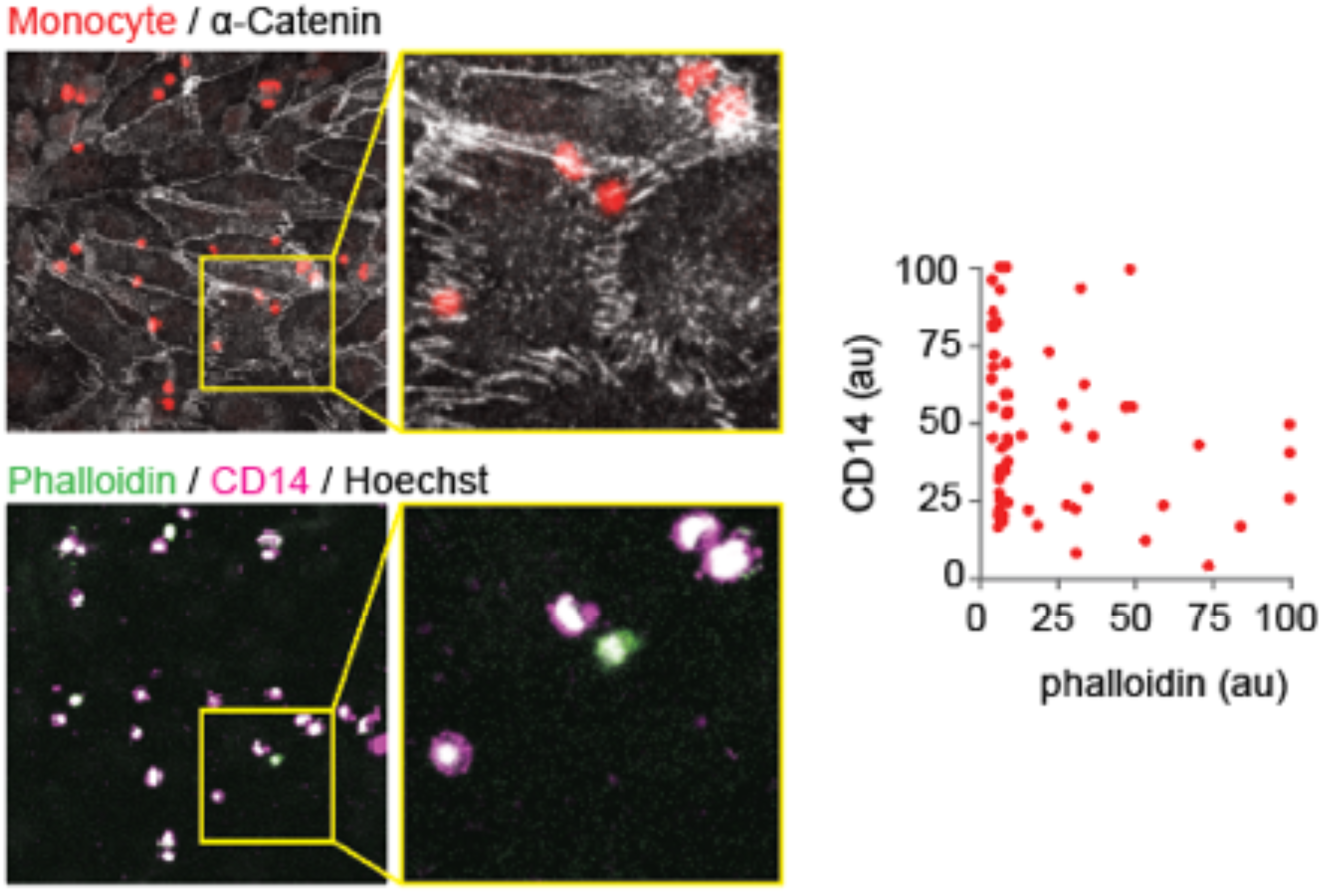
Immuno-cytochemistry indicates presence of classical and non-classical human monocytes on top of human umbilical vein cell layer. Cells were co-cultured, subsequently fixed and stained with markers for actin (green), Hoechst (blue) and CD14 (magenta). Note cells with low CD14 and high Phalloidin levels. Scale bars, 100 µm.

### SUPPLEMENTARY MOVIE LEGENDS

**Movie S1:**
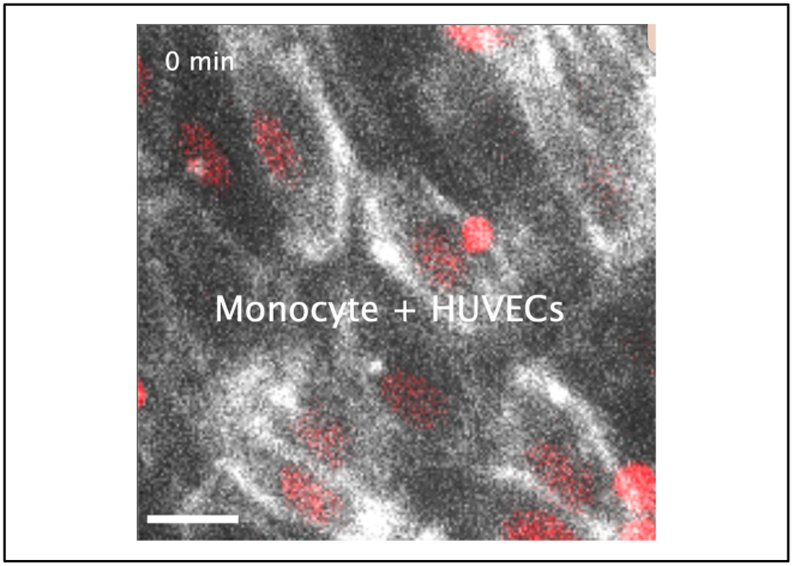
Motion pattern of primary human monocytes/macrophages on top of HUVECs (track length, 76 frames; speed, 10 fps, 1 frame per minute). Scale bar, 20 µm.

**Movie S2:**
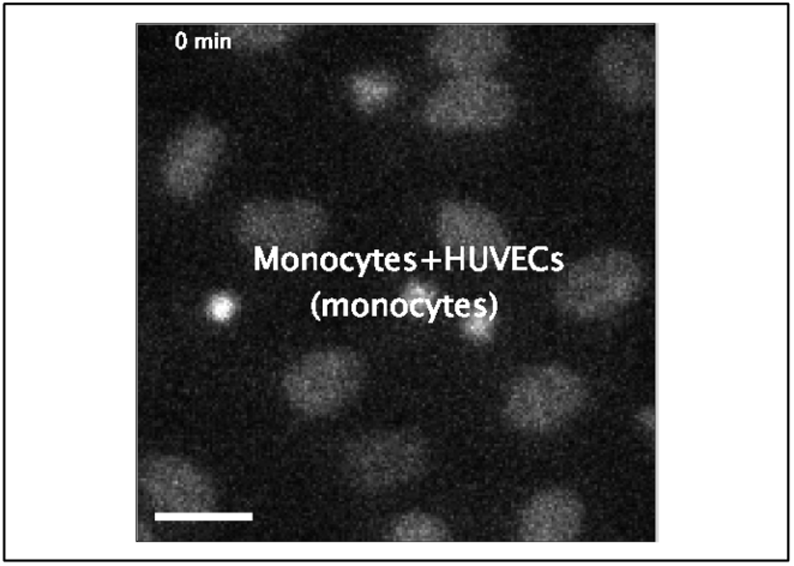
Motion pattern of primary human monocytes/macrophages co-cultured with HUVEC (track length, 91 frames; speed, 20 fps, 1 frame per minute). Scale bar, 20 µm.

**Movie S3:**
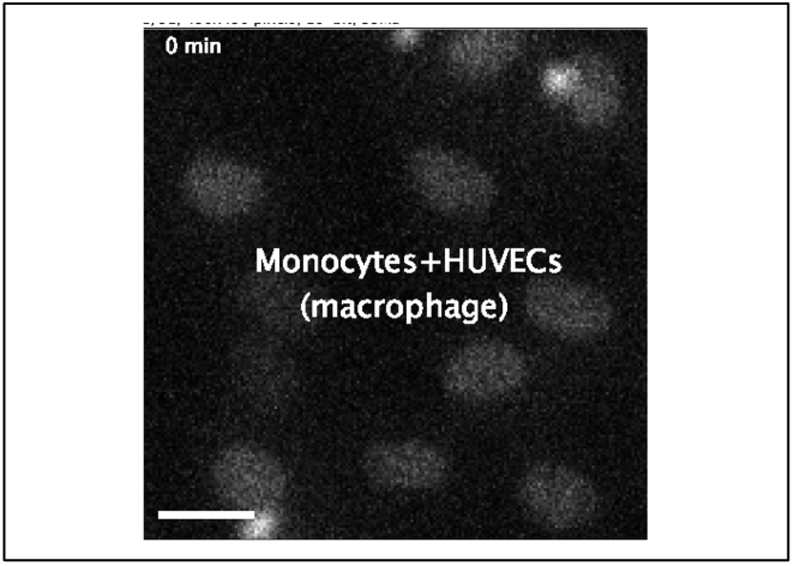
Motion pattern of primary human monocytes/macrophages co-cultured with HUVEC (track length, 121 frames; speed, 20 fps, 1 frame per minute). Scale bar, 20 µm.

**Movie S4:**
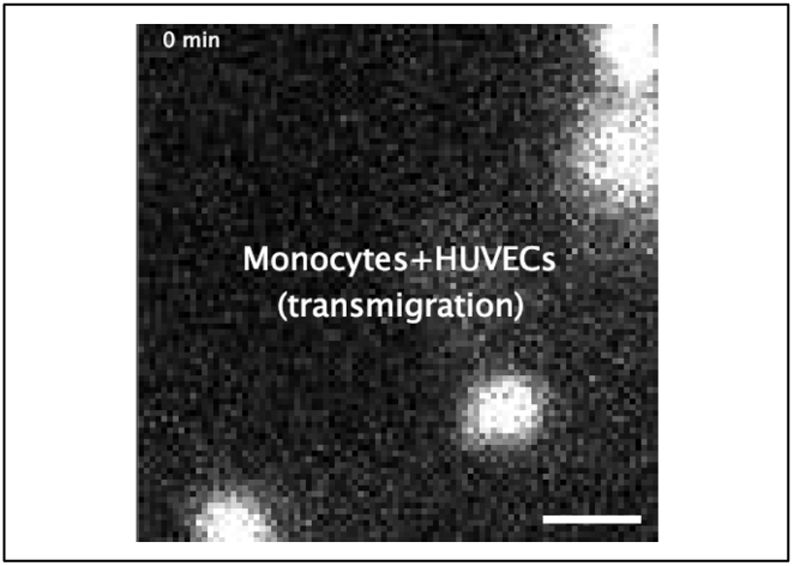
Transmigration of primary human monocytes/macrophages through HUVEC monolayer (track length, 32 frames; speed, 10 fps, 1 frame per minute). Scale bar, 10 µm.

**Movie S5:**
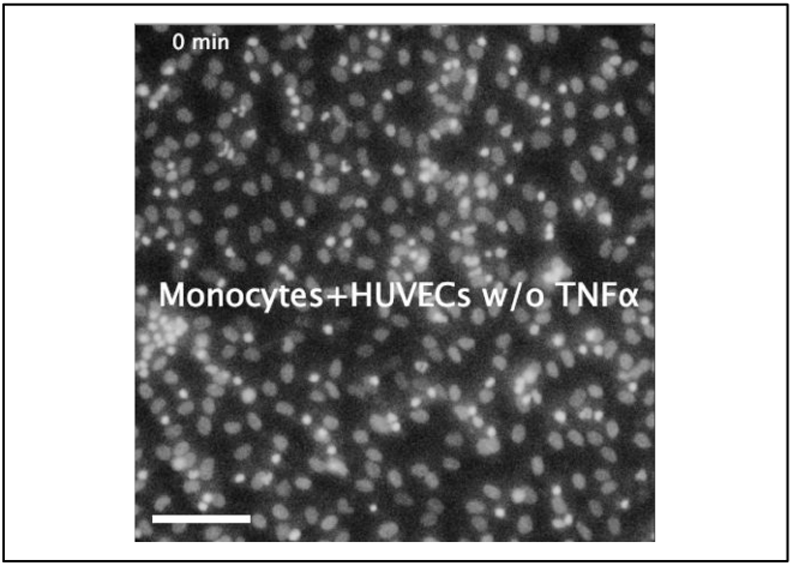
Motion pattern of primary human monocytes/macrophages on top of HUVECs (track length, 167 frames; speed, 20 fps, 1 frame per minute). Scale bar, 100 µm.

**Movie S6:**
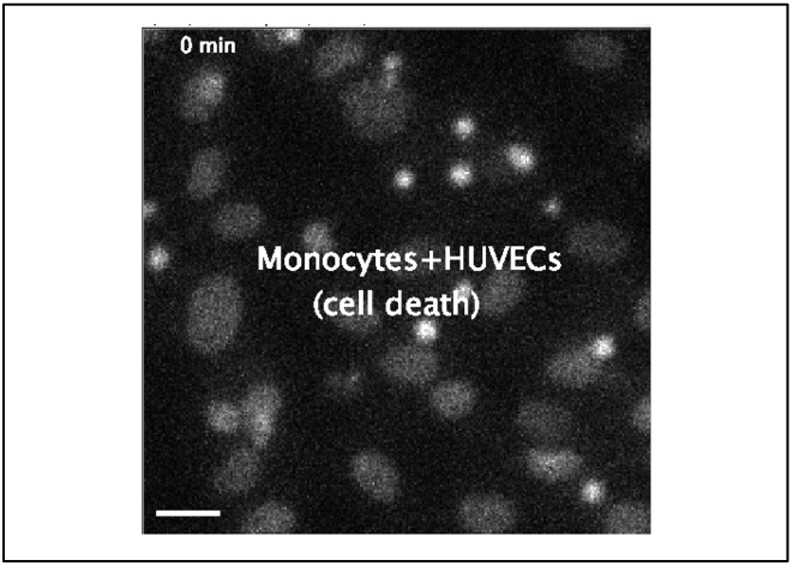
Motion pattern of primary human monocytes/macrophages in presence of dying cell. (track length 121 frames; 20 fps, 1 frame per minute). Scale bar, 20 µm.

**Movie S7:**
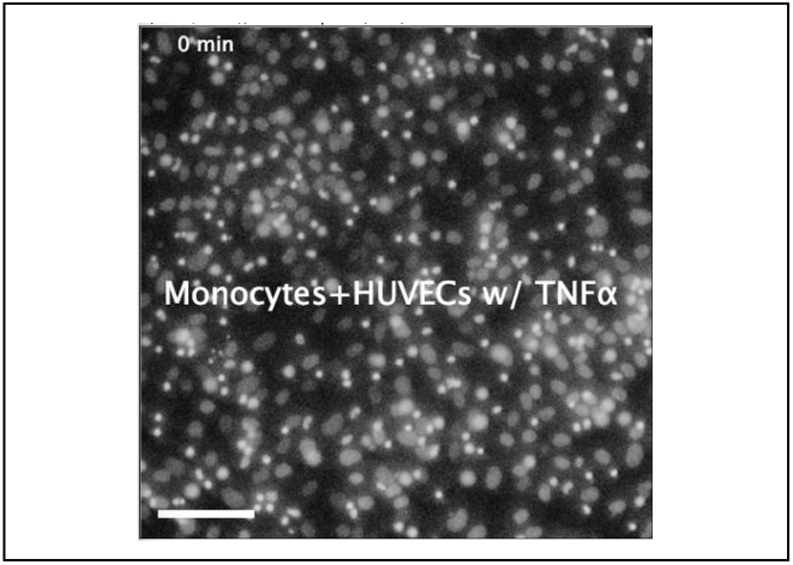
Motion pattern of primary human monocytes/macrophages on top of HUVECs in the presence of TNFα (track length 167 frames; speed, 20 fps, 1 frame per minute). Scale bar, 100 µm.

**Movie S8:**
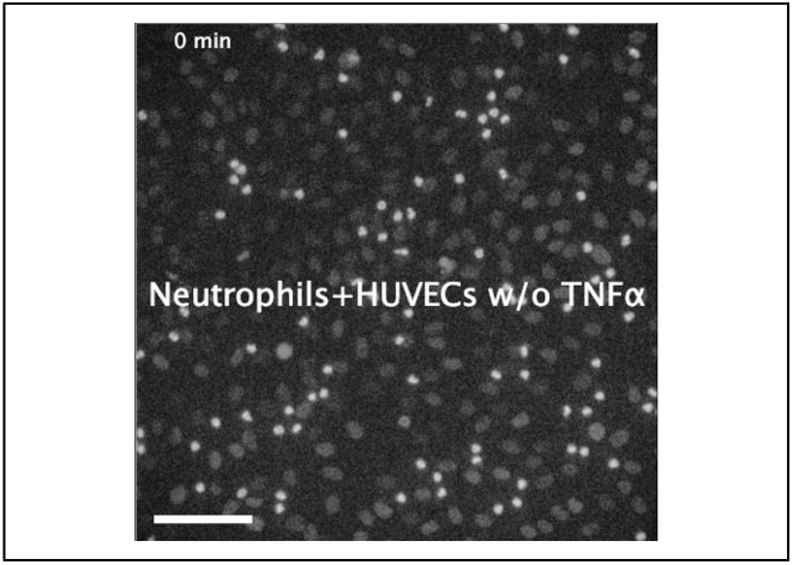
Motion pattern of primary human neutrophils on top of HUVECs (track length 167 frames; speed, 20 fps,2 frame per minute). Scale bar, 100 µm.

**Movie S9:**
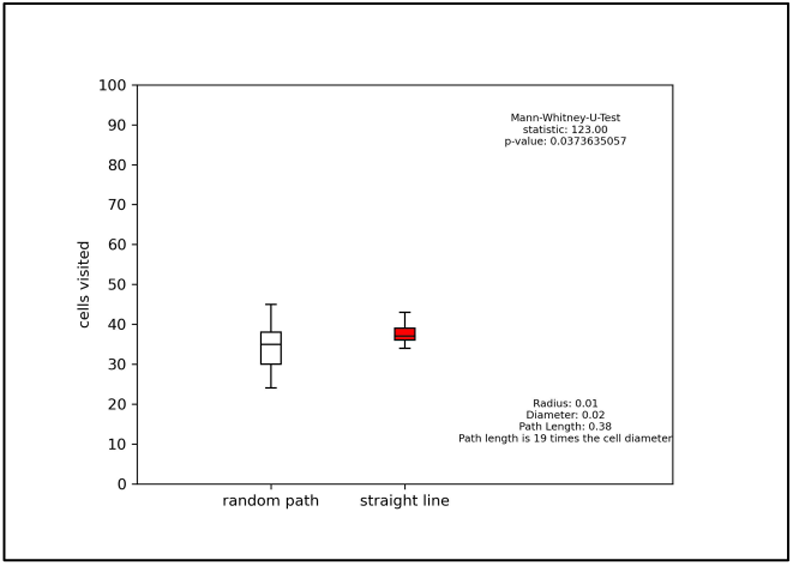
Numerical model depicting the number of sampled cells as function of track length for migration along cell boundaries (white, left) vs. straight line (red, right) (track length 40 frames; speed, 20 fps).

